# Mg^2+^-binding shifts the IM30 activity from membrane protection to membrane destabilization

**DOI:** 10.1101/2020.09.25.313916

**Authors:** Benedikt Junglas, Amelie Axt, Carmen Siebenaller, Hilal Sonel, Nadja Hellmann, Stefan A.L. Weber, Dirk Schneider

## Abstract

The inner membrane-associated protein of 30 kDa (IM30) is essential in chloroplasts and cyanobacteria. The spatio-temporal cellular localization of the protein appears to be highly dynamic and triggered by internal as well as external stimuli, mainly light intensity. A soluble fraction of the protein is localized in the cyanobacterial cytoplasm or the chloroplast stroma, respectively. Additionally, the protein attaches to the thylakoid membrane as well as to the chloroplast inner envelope or the cyanobacterial cytoplasmic membrane, respectively, especially under conditions of membrane stress. IM30 is involved in thylakoid membrane biogenesis and/or maintenance, where it either stabilizes membranes and/or triggers membrane-fusion processes. These apparently contradicting processes have to be tightly controlled and separated spatiotemporally in chloroplasts and cyanobacteria. The latter process depends on Mg^2+^-binding to IM30; yet, it still is unclear how Mg^2+^-loaded IM30 interacts with membranes and promotes membrane fusion. Here we show that interaction of Mg^2+^ with IM30 results in increased binding of IM30 to native as well as model membranes. Via Atomic Force Microscopy in liquid, IM30-induced bilayer defects were observed in solid-supported bilayers in presence of Mg^2+^. The observed interaction of IM30 with membrane surfaces differs dramatically from previously observed membrane-stabilizing, carpet-like structures in the absence of Mg^2+^. Mg^2+^-induced alterations of the IM30 structure switches the IM30 activity from a membrane-stabilizing to a membrane-destabilizing function, a crucial step in membrane fusion.

---

IM30, the *inner membrane-associated protein of 30 kDa*, also known as the *vesicle inducing protein in plastids 1* (Vipp1), is conserved in nearly all oxygenic photosynthetic organisms (1). The protein is a member of the PspA/IM30 family and likely has evolved via gene duplication from the bacterial *phage shock protein A* (PspA) (2).

Soluble IM30 is localized in the chloroplast stroma or the cyanobacterial cytoplasm, respectively. However, a fraction of the protein is attached to the chloroplasts’ inner envelope or the cyanobacterial cytoplasmic membrane, as well as to thylakoid membranes (TMs) (4–7). The activity of IM30 is broadly linked to the biogenesis/maintenance of TMs (recently reviewed in (8)), and two specific activities are currently studied and discussed in greater detail: (i) membrane protection (9–15) and (ii) membrane remodeling and repair (10, 15, 16). Protection and maintenance of the TM system are necessary to prevent energy dissipation and to maintain a proton gradient across the TM, a membrane system highly vulnerable to membrane damage (17). IM30 appears to share its membrane-stabilizing and/or protecting activity with other proteins of the PspA/IM30 family, namely PspA and LiaH (reviewed in (18)). Membrane protection via IM30 and related proteins likely involves the formation of a protecting protein carpet on a membrane surface (19) to block proton leakage of damaged membranes (15). Remodeling of the internal TM system, on the other hand, is unique to cyanobacteria and chloroplasts where it is essential to adapt the size and structure of the TM system during TM development and in response to changing light conditions (20–22). Yet, both membrane protection and remodeling of the TM are crucial for efficient photosynthesis. However, membrane stabilization and membrane fusion contradict each other, as membrane fusion processes typically involve (at least partial) membrane destabilization. Thus, the two experimentally observed processes have to be tightly controlled and separated spatiotemporally in chloroplasts and cyanobacteria (23, 24).

The IM30 monomer is predicted to be mainly α-helical with seven predicted α-helices (6, 25). The monomer has an intrinsic propensity to form large homo-oligomeric complexes of variable sizes (>1.5 MDa; ∼24 – 40 nm diameter; ∼14 – 15 nm height) with a ring-like shape and a pronounced spike structure (6, 25–29). Besides ring-shaped complexes, larger double rings and rod-shaped supercomplexes have also been observed with diameters and rotational symmetries similar to the ring complexes (6, 26, 28–30). However, the molecular structure of these complexes, as well as the monomer structure is still unknown (9, 26). Yet, while IM30 monomers appear to be largely α-helical when incorporated into rings (as predicted in (6)), a recent study demonstrated that a large part of IM30 is intrinsically disordered when the protein is monomeric or organized in smaller oligomers (19).

IM30 preferentially binds to negatively charged membrane surfaces, which changes the order and polarity in the lipid headgroup region (9, 10). Upon binding, IM30 rings disassemble, resulting in the formation of an IM30 carpet on the membrane surface that covers parts of the membrane, as recently shown *in vitro* (19). These carpets stabilize damaged membranes and thereby likely preserve the barrier function of the TMs (19). The *in vitro* observations explain the *in vivo* observed formation of large IM30 assemblies at defined TM regions in response to membrane stress conditions (23, 31). Yet, while the membrane protecting activity of IM30 has been observed, IM30 triggers membrane fusion when Mg^2+^ is present (10). Notably, in chloroplasts and cyanobacteria, Mg^2+^ flux across the thylakoid membrane is used to counterbalance the light-induced formation of a proton gradient across the TM (recently reviewed in (32)). Once Mg^2+^ has entered the chloroplast stroma or cyanobacterial cytoplasm, respectively, out of the thylakoid lumen, it allosterically regulates the activity of several enzymes and proteins (33, 34). Indeed, Mg^2+^ directly binds also to IM30 in solution, inducing several structural rearrangements, resulting in an increased surface hydrophobicity and increased protease resistance of the protein’s C-terminal domain (35). Thus, IM30-triggered membrane remodeling is potentially controlled via Mg^2+^-binding to IM30. However, while recently the organization of membrane-protecting IM30 species on the membrane surface has been elucidated (19), critical steps of the membrane fusion/remodeling process are still unclear. How does Mg^2+^-binding to IM30 affect the interaction of the protein with membranes, the membrane structure and induces membrane fusion?

Here we show that IM30 binds better to native as well as to model membranes when Mg^2+^ is present. Importantly, membrane adsorption onto solid-supported bilayers substantially differs in the presence of Mg^2+^ from membrane adsorption in the absence of Mg^2+^, as shown here via Atomic Force Microscopy: Upon binding of Mg^2+^-loaded IM30 (short: IM30(Mg^2+^)) to membrane surfaces, we observed the emergence of membrane defects in close proximity to the IM30 binding region. Membrane destabilization by IM30(Mg^2+^) and the formation of membrane defects, as observed in the present study, now explain the recently observed membrane fusogenic activity of IM30. Thus, we conclude that IM30 has two (seemingly opposing) functions, and Mg^2+^-binding to IM30 switches the protein function from “membrane protection” to a “membrane fusion” activity

## Results

### Mg^2+^ increases the membrane-binding propensity of IM30

Binding of Mg^2+^ to IM30 alters the structure of IM30 and triggers membrane fusion (10, 35). Due to the exposure of an increased hydrophobic surface upon Mg^2+^-binding (35), the structural changes induced by Mg^2+^-binding potentially enhance its membrane binding affinity. To test this assumption, we first assessed whether binding of IM30 to native TMs depends on the cytosolic Mg^2+^ concentrations. To address this, we determined the amount of membrane-bound IM30 in the presence of different Mg^2+^ concentrations using cellular extracts of the cyanobacterium *Synechocystis* sp. PCC 6803 (from here on *Synechocystis*) via sucrose density gradient centrifugation and subsequent immuno-detection of membrane-attached IM30 (Figure 1A). When increasing concentrations of Mg^2+^ were added to the *Synechocystis* cell lysate, an increasing amount of native IM30 was found to colocalize with membranes in the sucrose gradient, strongly indicating increased membrane binding (Fig. 1A). In contrast, the amount of IM30 colocalizing with the TM in a control, where endogenous Mg^2+^ was removed via EDTA, was below the detection limit of our immunoblots. Thus, Mg^2+^-binding to IM30 clearly enhances its ability to interact with cyanobacterial membranes.

**Figure 1:**
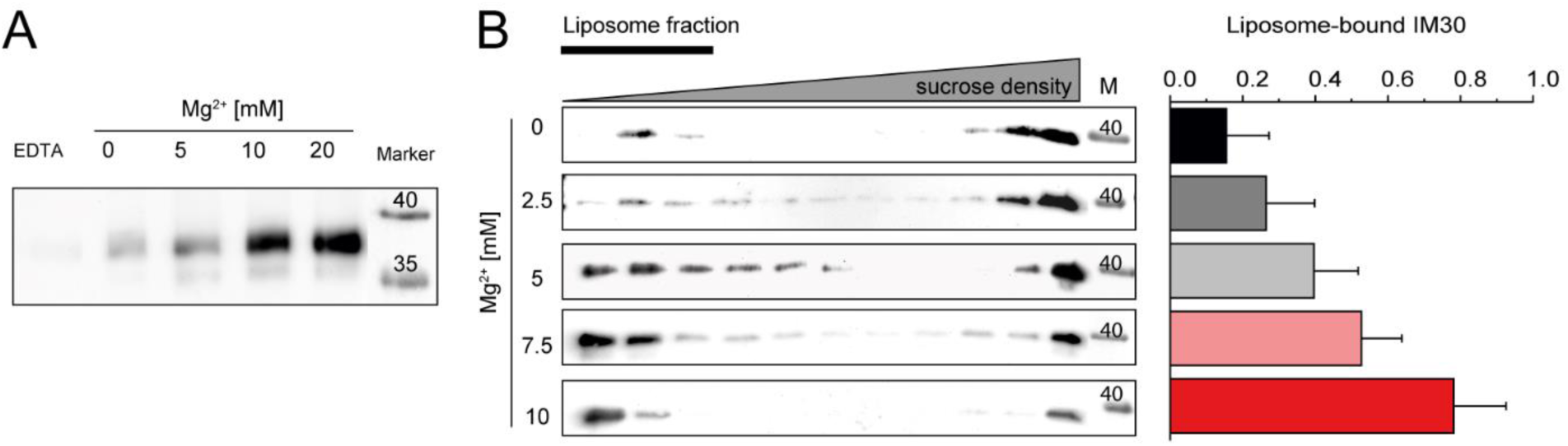
The membrane binding propensity of IM30 is increased in the presence of Mg^2+^. **A:** The amount of IM30 bound to TMs in the presence of increasing Mg^2+^ concentrations was estimated via sucrose density gradient centrifugation of *Synechocystis* cell lysate and subsequent immunoblotting of the TM fraction. Endogenous IM30 were not detected to colocalize with TMs when 20 mM EDTA was present to remove any endogeneous Mg^2+^. At concentrations of 5 mM Mg^2+^ or higher, increasing amounts of IM30 could be detected. **B:** Interaction of IM30 with 100% DOPG liposomes in presence of 0 to 10 mM Mg^2+^ was detected via sucrose density gradient centrifugation. Localization of IM30 in the density gradient was monitored via immunoblotting, and liposomes were localized using the incorporated fluorescent lipid dye NBD-PE (Suppl. Fig S1). The amount of liposome-bound IM30 was quantified via densitometric analysis of the immunoblot bands (bar chart). An increasing amount of IM30 colocalizes with the liposomes with increasing concentrations of Mg^2+^.

To test whether IM30 binds directly to membranes rather than to exposed proteins or protein domains that were copurified with the *Synechocystis* membrane, we next used negatively charged model membranes (100 % DOPG liposomes). We performed sucrose density gradient centrifugation and subsequent immunological detection of the protein at increasing Mg^2+^ concentrations (0-10 mM) (Fig. 1B). As shown before (10), we observed colocalization of a small IM30 fraction with liposomes in the absence of Mg^2+^, indicating low-affinity binding of IM30 to liposomes in absence of Mg^2+^. Based on the immunological analyses (Fig. 1B), only ∼15% of IM30 colocalized with the liposomes (fraction 1 to 3, Suppl Fig. S1), whereas the majority of the protein was found in the pellet fraction, as the high molecular mass IM30 assemblies sedimented during centrifugation. However, increasing the Mg^2+^ concentration to up to 10 mM largely increased the liposome-bound fraction, finally resulting in ∼80% bound IM30 at 10 mM Mg^2+^ (Fig. 1B).

Next, we directly visualized binding of IM30 to lipid membranes using DOPG/DOPC (20/80) giant unilamellar vesicles (GUV) in the absence and presence of Mg^2+^ via fluorescence microscopy (Fig. 2). We used recombinantly produced and purified IM30 C-terminally fused to CFP (IM30-CFP), and the GUVs were labeled by doping the membranes with Atto633-PE. In absence of Mg^2+^, binding of the protein to the GUVs was not detectable (Fig. 2). However, upon addition of 5 mM Mg^2+^, binding of IM30-CFP to the GUVs was clearly visible, and the CFP-fluorescence was evenly distributed over the GUV area (Fig. 2). Note that the GUVs were highly unstable at higher concentrations of negatively charged lipids and/or >5 mM Mg^2+^ concentrations. Thus, we were limited to lower ratios of PG in the GUVs and lower Mg^2+^ concentrations compared to the centrifugation assay. Nevertheless, the amount of IM30 bound to negatively charged GUVs was also clearly increased in the presence of Mg^2+^.

**Figure 2:**
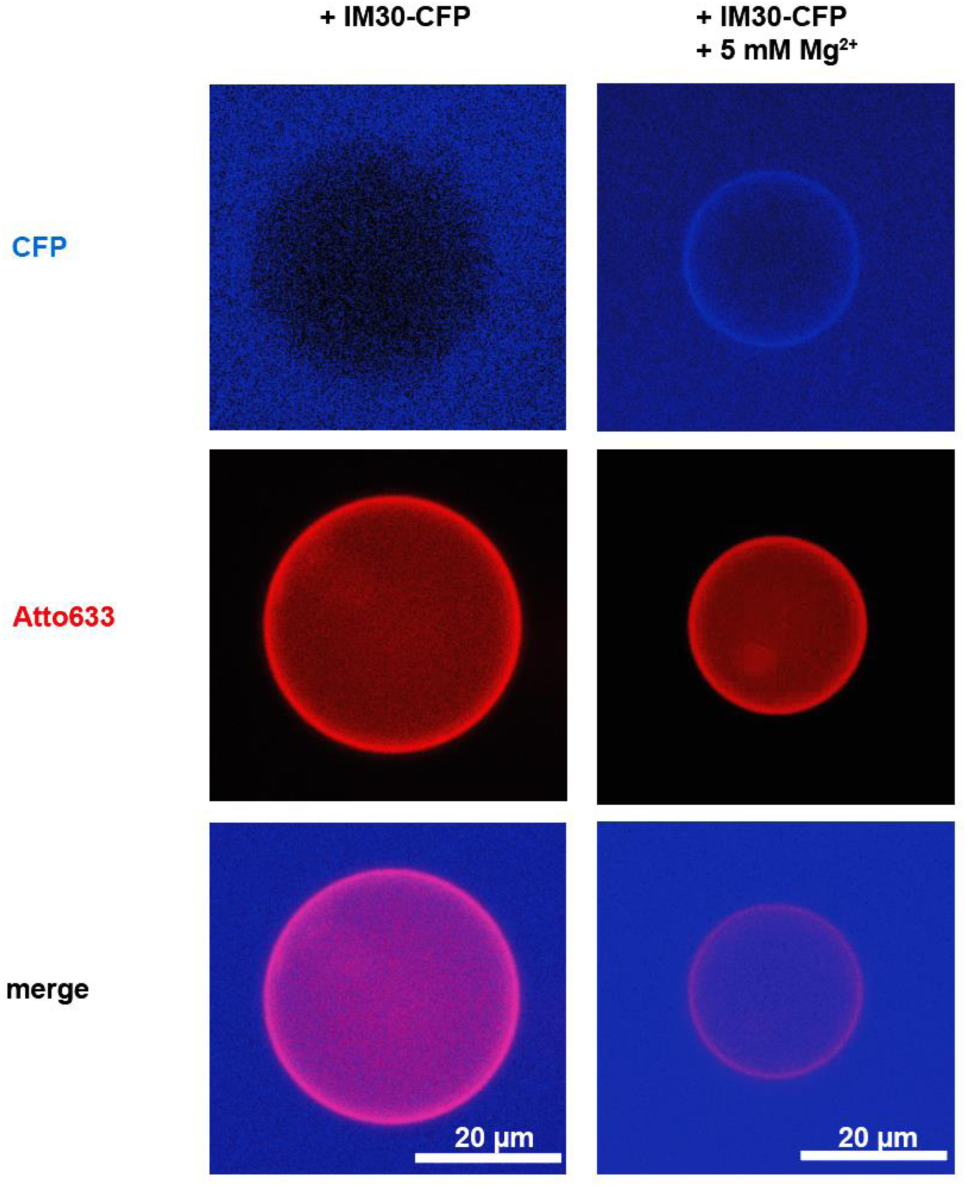
Increasing amounts of IM30 bind to GUVs in presence of Mg^2+^. Adsorption of IM30-CFP to DOPG/DOPC (20/80) GUVs in absences *vs* presence of Mg^2+^ was visualized using fluorescence microscopy. GUVs were detected by the fluorescence signal of the incorporated lipid dye Atto-633-PE. IM30-CFP was visualized via monitoring the CFP fluorescence. The amount of adsorbed IM30 was increased at 5 mM Mg^2+^ (B) when compared to Mg^2+^-free buffer (A), as indicated by the increased CFP fluorescence intensity localized on the GUV surface.

Thus, Mg^2+^-binding to IM30 clearly boosts its ability to bind to native as well as to model membranes.

### Mg^2+^ binding alters the viscoelastic properties of IM30 decorated membranes

To quantitatively analyze the impact of Mg^2+^ on IM30 membrane binding, we next used a quartz-crystal microbalance (QCM) to follow the binding of IM30 to a solid supported lipid bilayer (SLB) (20% DOPG / 80% DOPC), as previously described in the absence of Mg^2+^ (15). In QCM experiments, the resonance frequency of a quartz crystal is determined. This resonance changes when additional material binds to the crystal. Hence, binding of any material, *e.g*. IM30, to the SLB (which is attached to the quartz crystal) leads to a frequency shift towards lower resonance frequencies. Assuming that the adsorbed material is rigid, the total mass bound to the crystal can be determined via the Sauerbrey equation (36, 37). However, when a viscoelastic layer forms on the crystal, the oscillation is damped, leading to a decreased frequency shift, and the mass determined based on the Sauerbrey equation then underestimates the mass actually adsorbed. The Sauerbrey equation is applicable only when the damping signal in Hz is much smaller than the frequency shift. Yet, it is possible to obtain information on the viscoelastic properties of adsorbed material via analyzing the damping/frequency shift ratio (38). Thus, we were able to determine changes in IM30 surface binding and to obtained information about the viscoelastic properties of the membrane-bound IM30.

As can be seen in Figure 3A, binding of IM30 to the SLB caused a shift of the resonance frequency, associated with increased damping of the resonance oscillation in the absence of Mg^2+^. The frequency shift reached a constant level at about -80 Hz after ∼2500 s, as already shown previously (15). In stark contrast, in presence of Mg^2+^ the frequency signal changes were only ∼25% of the signal change observed in the absence of Mg^2+^ (Fig. 3A). However, in both cases the damping signal was large enough to indicate that the protein film is not rigid but viscoelastic. The reduced frequency shift in presence of Mg^2+^ could be caused by a lower amount of mass attached to the membrane surface and/or changes in the viscoelastic properties. In fact, the viscoelastic properties appear to have a higher contribution to the oscillation in presence than in absence of Mg^2+^, since the ΔΓ/Δf ratio is larger. In absence of Mg^2+^, the contribution of viscoelastic properties to the oscillation actually appear to get smaller, as indicated by the different slopes when the damping is plotted *vs* the frequency shift (Fig. 3B/C). This observation is in agreement with the observation that initially IM30 rings bind to the membrane which then convert into flat carpet-like structures (19). The mass increase upon ring binding is given by the sum of the protein mass and adhered water, in particular inside the ring. Upon ring disassembly this water is released. Furthermore, the contribution of elastic and viscous behavior might shift in this process. The viscoelastic properties observed in presence of Mg^2+^, which stabilizes the ring structures (35), are more similar to the viscoelastic properties observed in the initial phase of IM30 membrane-interaction in the absence of Mg^2+^ (Fig. 3C). In summary, the QCM measurements clearly indicate different interaction of IM30 with membrane surfaces in absence *vs* presence of Mg^2+^.

**Figure 3:**
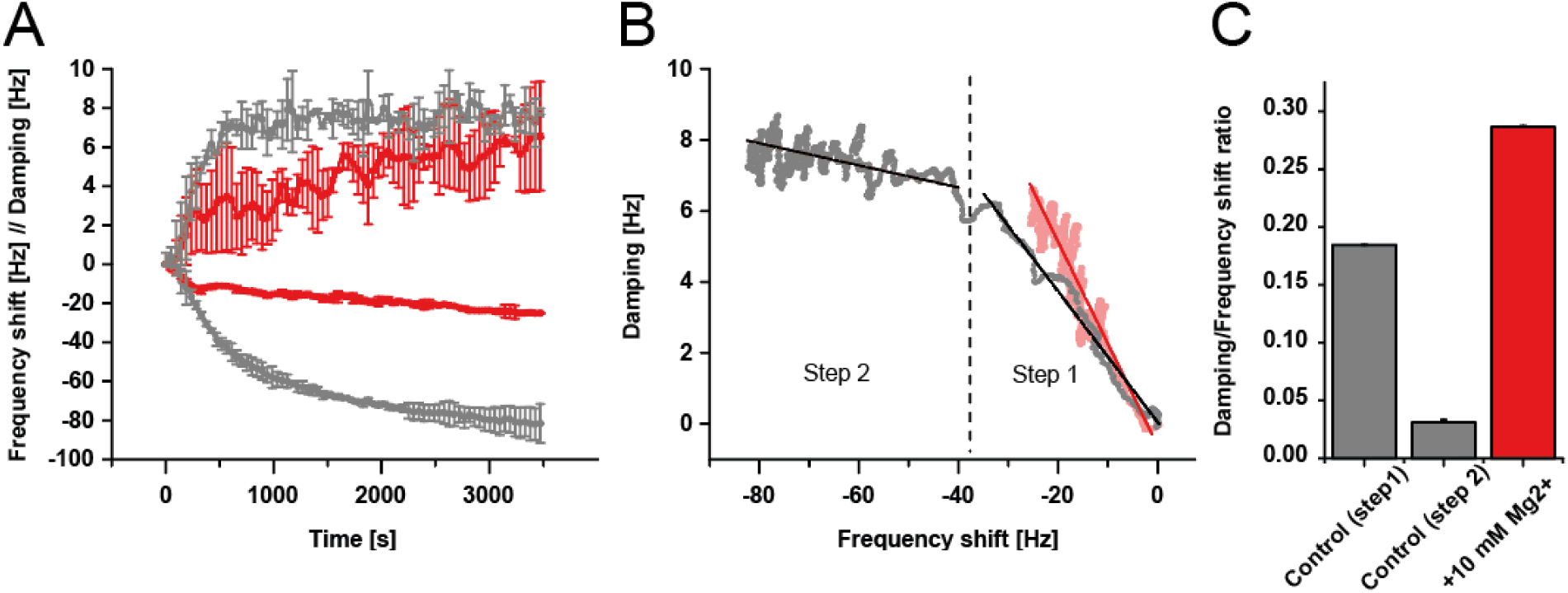
The viscoelastic properties of membrane-bound IM30 differ in presence and absence of Mg^2+^. **A:** The frequency shift (negative values) and damping (in terms of the resonance peak’s full width at half maximum; positive values) of QCM measurements of IM30 binding to an SLB (80% DOPC/20% DOPG) in the absence (dark grey, (15) and presence (red line) of 10 mM Mg^2+^ are shown. IM30 binding to the SLB in the presence of Mg^2+^ caused a smaller frequency shift than in the absence of Mg^2+^. The damping signals reached about the same levels. Error bars represent SD, n=3. For a better overview, only every 30th data point is shown. **B:** A plot of the frequency shift *vs* the damping of IM30 binding to an SLB reveals a shift in viscoelastic properties during protein binding in the absence of Mg^2+^ (red) and a single phase in the presence of Mg^2+^ (red). The regimes were determined by the slopes of linear fits: Step 1 in the absence of Mg^2+^: m=-0.1844±0.0002; (R^2^=0.99959); Step 2 in the absence of Mg^2+^: m=-0.031±0.002; (R^2^=0.08976); Step 1 in the presence of Mg^2+^: m=-0.28742±0.0008; (R^2^=0.98384). Error bars are not shown, n=3. **C:** A comparison of the damping/frequency shift ratio in the absence and presence of Mg^2+^ is shown.

### Mg^2+^ triggers IM30 induced formation of bilayer defects

To better understand how IM30 affects the membrane structure when Mg^2+^ is present, and *vice versa*, we next analyzed membrane adhesion of IM30 on a nanometer-scale using AFM on SLBs. All AFM measurements were performed using 100% DOPG SLB on mica. Due to technical limitations, the first image after injection of IM30 could only be scanned after at least 10 minutes. Therefore, processes happening in the first 10 minutes after injection were not observed.

After membrane adhesion, IM30 formed particles of variable heights and diameters on the bilayer in the absence as well as in the presence of Mg^2+^, but the dimensions of the particles differed (Fig. 4 A/B). Therefore, we at first performed a statistical analysis of the particle size, only taking into account particles that were larger than one pixel and extracted the particle properties with the AFM analysis software GWYDDION (Fig. 4). In absence of Mg^2+^, IM30 particles have a mean diameter of 38 nm and a mean height of 0.8 nm (Fig. 4 C). 67 % of the particles have diameters in the range of ∼20 nm to 40 nm. These diameters are in line with diameters of IM30 rings determined via TEM (∼20 – 35 nm (26)) as well as IM30 rings bound directly to the *mica* surface and visualized via AFM (outer diameter ∼30 – 60 nm (19)). Yet, the height of the particles is not in line with previous findings: estimations based on negative stain TEM reconstructions suggest a height of 13-15 nm for the rings (26), and recent AFM studies in liquid environments report a height of 15-25 nm for IM30 rings attached to *mica* surfaces (19). The low height of the particles, together with the slightly increased diameters, potentially indicates beginning or partial disassembly of IM30 ring structures on the membrane surface. This was also indicated by our QCM measurements (Fig. 3B/C) and is in line with the finding that in the absence of Mg^2+^ IM30 rings disassemble upon membrane binding and rearrange into large and flat membrane protecting carpets upon prolonged interaction with PG bilayers (see Fig. S2 and (19)).

**Figure 4:**
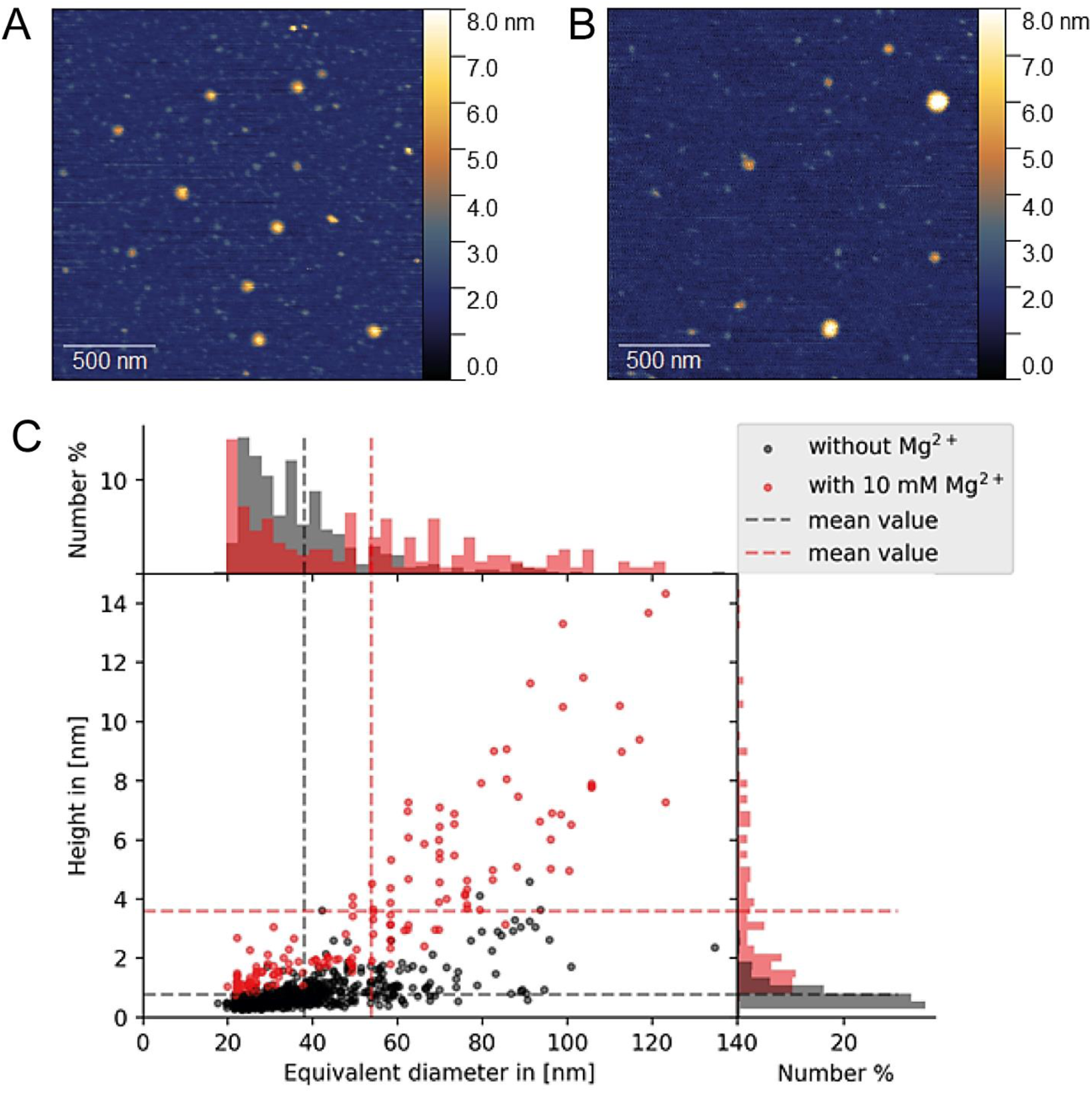
Dimensions of membrane-bound IM30 particles. **A:** Example for AFM topography image used for particle analysis showing IM30 on a DOPG SLB in the absence of Mg^2+^. Heights are indicated in the image using a false-color ruler. **B:** IM30 on a DOPG SLB is shown in the presence of Mg^2+^. Heights are indicated in the image using a false-color ruler. **C:** The diameter plotted *vs* the height of IM30 *punctae* on the DOPG SLB in the absence (black n=614), and the presence of 10 mM Mg^2+^ (red, n=156) were determined using GWYDDION particle analysis (49). The particle count in the absence of Mg^2+^ is higher since these measurements were more stable, and we were able to scan more surface area per measurement. The normalized histograms on the sides indicate the distribution of the particle dimensions. The dashed lines indicate the mean particle dimensions. Mean particle diameter in absence of Mg^2+^: 38 nm *vs* 53.7 nm in presence of Mg^2+^; mean particle height in absence of Mg^2+^: 0.8 nm *vs* 3.6 nm in presence of Mg^2+^.

In stark contrast, when membrane binding of IM30 was analyzed via AFM in presence of 10 mM Mg^2+^, the mean particle diameter is 53.7 nm and the distribution of diameters is very broad (40 % have diameters from 20 to 40 nm, 47 % from 40 to 90 nm and 13 % have diameters above 90 nm) (Fig. 4). Nevertheless, 60 % of the particles have larger diameters than the mean diameter (38 nm) of the particles observed in the absence of Mg^2+^ (compare Fig. 4). The mean height of the particles in the presence of Mg^2+^ is 3.6 nm, while 44% of particles were below 2 nm in height (Fig. 4 C). The determined height is also smaller than the height of intact rings (26), again indicating ring rearrangement and/or disassembly on the membrane surface. However, our statistical analysis indicates that in the presence of Mg^2+^, 99 % of the particles are higher than the mean height of particles in absence of Mg^2+^. In addition, the height distribution of the particles is much more heterogeneous in presence of Mg^2+^: 37% have heights between 1 and 2 nm, 43 % have heights from 2 – 7 nm, and 13 % have heights above 7 nm. In contrast, without Mg^2+^ the height does not exceed 4.6 nm and increases only minimally with increasing particle width. Hence, the particle diameters differ significantly in absence *vs* presence of Mg^2+^. In fact, in presence of Mg^2+^ IM30 rings are stabilized and increased ring-stacking and formation of double rings was observed in solution (35). The increased height of membrane-attached IM30 in presence of Mg^2+^ likely originates from stabilization of IM30 supercomplexes and membrane attachment of higher-ordered IM30 structures and explains the higher viscoelasticity observed in the QCM measurements (Fig. 3B/C).

In contrast to the situation in absence of Mg^2+^ (19), we did not observe the formation of carpet structures on the membrane surface when Mg^2+^ was present. Instead, in presence of Mg^2+^, individual *punctae* started to flatten out over time and seem to initiate formation of a lipid bilayer defect (Fig. 5). These defects grew in size and depth and ranged from 10 - 36 nm diameter pores to larger defects spanning several hundred nm of the SLB. Some *punctae* started to form pores shortly after their appearance on the bilayer (∼20–30 min), whereas other *punctae* converted into pores only after prolonged incubation times (1 – 2 h) and others remained stable over the whole experiment (5 – 6 h). (Only *punctae* that did not show signs of defect formation were taken into account for the statistical analysis shown in Fig. 4).

**Figure 5:**
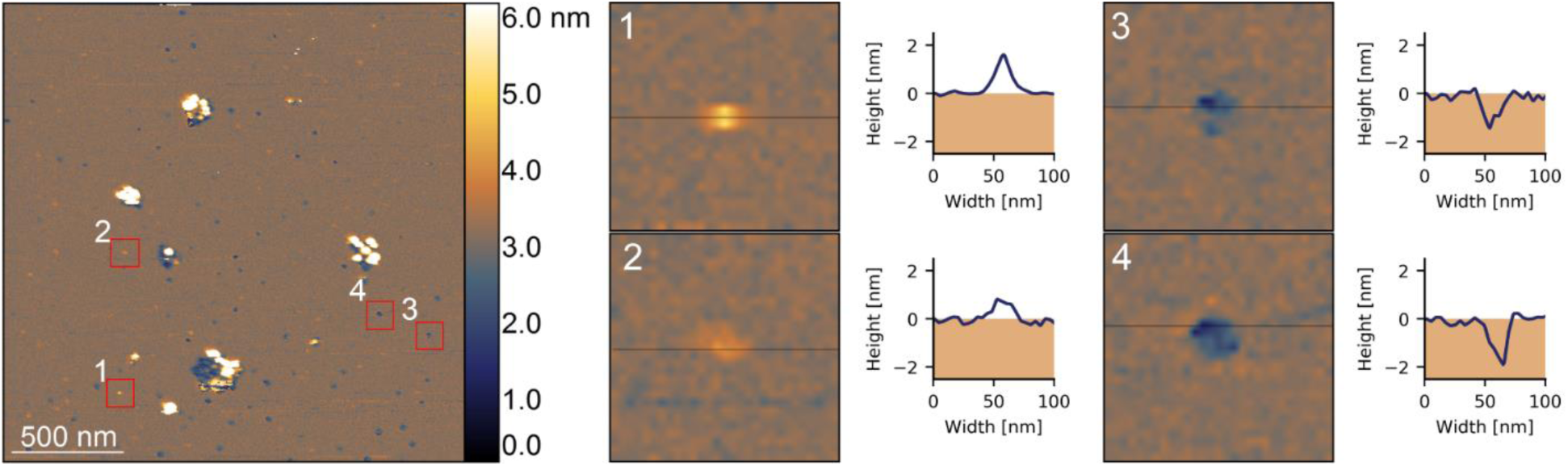
IM30 generates bilayer defects in the presence of Mg^2+^. AFM topography images of IM30 on a DOPG SLB in the presence of Mg^2+^ are shown together with false-color rulers indicating the height. Profile lines indicate the position of the cropped images shown on the right. On the right, a series of details are shown with their corresponding height-profiles, illustrating representative steps of IM30-mediated formation of bilayer defects.

**Figure 6:**
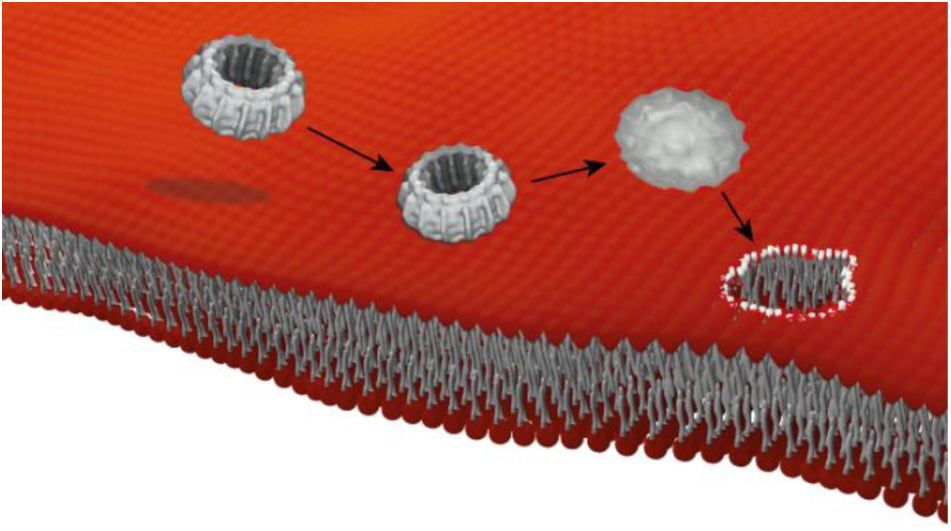
Bilayer defect formation induced by IM30 binding. Upon membrane binding, IM30 rings likely disassemble to a flat unordered assembly of small oligomers and a membrane pore starts to form that is likely stabilized by small IM30 oligomers at the bilayer edges. The models were created using BLENDER and CHIMERAX (50).

## Discussion

Previously, it has been shown that IM30 forms carpet structures on negatively charged membrane surfaces that can stabilize membranes (19). This observation is in perfect agreement with a membrane protecting activity suggested before for IM30 and PspA family members, as well as with the formation of large membrane covering structures under (membrane) stress conditions observed *in vivo* (23, 31). Yet, IM30 has also been shown to mediate membrane fusion, at least *in vitro*, which involves membrane destabilization. However, as shown recently, the presence of Mg^2+^ is required for IM30 acting as a membrane fusion protein (10, 35).

Binding of IM30 to negatively charged membranes has been observed using small PG liposomes, PC/PG GUVs, and PG SLBs in the absence of Mg^2+^ (Fig. 1-4). Interestingly, membrane interaction of IM30 clearly is enhanced at increasing Mg^2+^ concentrations, where the amount of protein bound to PG containing membranes as well as to isolated cyanobacterial membranes increases (Fig. 1 and 2). Mg^2+^ directly binds to IM30 (35), resulting in a rearrangement of the IM30 structure, involving exposure of extended hydrophobic surface regions on IM30 rings, which promotes membrane binding (35). However, we currently cannot completely exclude further effects of Mg^2+^ binding to the negatively charged membrane surface, which could *e.g.* involve “bridging” the negatively charged lipid head groups and negatively charged residues at the protein surface. However, the structures of membrane-bound IM30 observed in the present study in the presence *vs* absence of Mg^2+^ clearly differ: In the presence of Mg^2+^ the adsorbed species seem to be more viscoelastic (Fig. 3C), and the small-diameter *punctae* initially forming in the presence of Mg^2+^ have an increased height when compared to *punctae* forming in the absence of Mg^2+^ (Fig 4). This likely reflects the overall increased stability and compactness of the IM30(Mg^2+^) protein (35). In particular after longer incubation times, the impact of Mg^2+^ on IM30 membrane adhesion and the membrane structure and stability became evident: While IM30 forms large membrane-stabilizing carpet structures on a DOPG membrane surface in the absence of Mg^2+^ (19) (Fig. S2), it induces localized bilayer defects in the presence of Mg^2+^. These defects start as small holes within individual IM30 *puncta* and expand to larger defect structures (Fig. 5). The formation of bilayer defects (Fig. 5) also contributes to the reduced frequency changes observed in the QCM measurements in the presence of Mg^2+^ (see Fig. 1). It is fair to assume that the processes observed in the AFM measurements took also place during the time course of the QCM measurement, and the local loss of lipid material lead to a reduced lipid-covered area of the quartz crystal, and consequently to a reduced frequency shift.

Formation of large membrane defects, as observed in presence of Mg^2+^, may be induced by initial binding of IM30(Mg^2+^) to and stabilizing spontaneously occurring, transient small bilayer defects. Sequential local accumulation of more IM30 protomers potentially promotes growth of the bilayer defects that might finally result in a bilayer-spanning toroidal or barrel-stave pore. Similar mechanisms for pore formation were suggested for membrane destabilization via antimicrobial peptides (AMPs) (39) and were already monitored using AFM (40). The irregularities in sizes and shapes of the pores support a toroidal pore model, where peptides pull lipid head groups inwards to line an inner pore wall and thus the resulting pores are lined with both peptides and lipids (41). However, also formation of bilayer defects via a detergent-like lipid solubilization mechanism are discussed for many AMPs (39, 42, 43), and it is not yet clear which is the prevailing mechanism.

However, it is also possible that membrane-bound IM30 actively induces, rather than recognizes, bilayer defects by the charge and surface tension imbalance that goes along with asymmetric bilayer binding of IM30, followed by bilayer rupture. Such an impact of asymmetric bilayer binding has been described for membrane remodeling intrinsically disordered proteins, which can effectively induce membrane curvature by steric/lateral pressure due to their larger steric volume compared to compact folded proteins (44). IM30 rings appear to disassemble upon membrane binding, and ring disassembly involves unfolding of the C-terminal half of the protein (18). It is well possible that IM30 locally bends membranes due to its asymmetry across the membrane: the soluble, intrinsically disordered IM30 domains are localized exclusively on the surface of one bilayer leaflet, and accumulation or larger protein domains at only one membrane face could result in increased steric repulsion on the membrane, inducing curvature and finally resulting in local membrane rupture.

### Conclusion

Mg^2+^ binding to IM30 results in an increased membrane binding propensity of IM30 as well as in an altered structure of membrane-bound IM30. While IM30 forms a membrane-protecting surface carpet in the absence of Mg^2+^, our observations now reveal that IM30 induces bilayer defects in the presence of Mg^2+^. The bilayer defects observed on SLBs likely trigger liposome fusion in solution. Consequently, IM30 has two (opposing) functions that are spatiotemporally controlled via Mg^2+^-binding. In photosynthetic organisms, at low Mg^2+^ concentration in the cytosol/lumen, IM30 might mainly fulfill its membrane chaperoning function. When the Mg^2+^ concentration increases (*e.g.* at increasing light intensity) IM30 becomes membrane fusion competent and eventually triggers membrane remodeling. Thus, Mg^2+^-binding switches from a membrane protecting to a membrane destabilizing IM30 activity.

## Experimental Procedures

### Cloning, expression, and purification of IM30

IM30 from *Synechocystis* PCC 6803 *sp.* was heterologously expressed in *Escherichia coli* BL21 DE3 and purified as described in detail previously (9, 10, 35, 45). In short, after expression, cells were lysed by sonification in lysis buffer (50 mM Na-phosphate, 20 mM imidazole, 300 mM NaCl, pH 7.6). The lysate was cleared by centrifugation, and the supernatant was applied to a Ni^2+^-NTA affinity column. The column was washed with increasing concentrations of imidazole, and the protein was eluted with 500 mM imidazole. The elution fractions were pooled and the buffer was changed to 20 mM HEPES pH 7.6 by gel-filtration (Sephadex G25).

### Liposome preparation

DOPG (*dioleoylphosphatidylglycerol;* Avanti Polar lipids) or a mixture of DOPC (*dioleoylphosphatidylcholine*; Avanti Polar lipids) and DOPG dissolved in chloroform was dried under a gentle stream of nitrogen for 10 min and under vacuum desiccation overnight. The resulting lipid film was rehydrated in 20 mM HEPES (pH 7.6) buffer to generate multilamellar liposomes. This liposome suspension was subjected to five freeze and thaw cycles resulting in the formation of unilamellar liposomes. If necessary, the liposomes were sonified in an ultrasonic bath cleaner to produce a more homogenous suspension (*e.g.* for AFM experiments) or they were sonified with a tip-sonicator in an ice bath followed by centrifugation (10 min 16500 g) to produce smaller liposomes (*e.g.* in QCM measurements). For the sucrose density gradient experiments, the liposomes were sized via 15 times extrusion trough a 50 nm filter, using an extruder from Avanti Polar lipids.

### Giant unilamellar vesicles

Giant unilamellar vesicles (GUVs) were generated by gel-assisted swelling of a lipid film on a dried PVA-film (*polyvinylalcohol*, MW 145000 Da; Merck Millipore). 30 µl of a 1% PVA-solution was heated to about 90 °C on a glass coverslip for 30 min. After cooling to RT, 3 µL of a 3 mM lipid solution (20% DOPG, 80% DOPC + 1:1000 Atto 633-DOPE, dissolved in chloroform) were spread on the PVA film and dried under a gentle stream of nitrogen. GUVs were formed by the addition of 200 µL of the corresponding swelling buffer. To analyze binding of IM30 to the GUVs, 20 mM HEPES buffer (pH 7.6) ±5 mM MgCl_2_ were used as swelling buffers. After 45 min, the GUVs were transferred to a new observation well, and 1 µM IM30-CFP was added carefully. GUVs were observed with a 20x objective using a Zeiss Axio Observer Z.1 fluorescence microscope with the Colibri 7 illumination module. The GUVs containing the Atto-633 fluorophor and the IM30-CFP proteins were detected using appropriate filters. The Zeiss ZEN software was used for image processing.

### Sucrose density gradients and IM30 binding

To analyze the binding of IM30 to DOPG liposomes at different Mg^2+^ concentrations, sucrose density gradient (SD) centrifugations were performed. All SD gradients were produced and fractionated using the Gradient Station (Biocomp). 160 µg DOPG liposomes (0.1 % NBD-DOPE) were incubated with 12 µg IM30 for 60 min at RT in 120 µL 20 mM HEPES, pH 7.6. The samples were loaded onto linear sucrose density gradients (5-50%) in 20 mM HEPES buffer (pH 7.6) containing 0, 2.5, 5, 7.5 or 10 mM MgCl_2_. Centrifugation was performed at 40000 rpm for 6 h at 25 °C followed by immediate fractionation. The individual fractions were analyzed by SDS-PAGE with subsequent immunoblot analysis to determine the protein distribution. An anti-IM30 antiserum (6) was used for analysis and antibody-binding was visualized via luminescence on a STELLA imaging system (Raytest). The immunoblots were analyzed quantitatively by using the densitometric analysis of the program ImageJ (46). The liposome distribution was analyzed via fluorescence spectroscopy detecting the lipid-coupled NBD fluorophore.

Binding of IM30 to the *Synechocystis* membranes was analyzed at varying Mg^2+^ concentrations by sucrose density centrifugation of *Synechocystis sp. PCC 6803* cell lysate. Cells were grown under standard conditions and lysed as described in (10, 45). The cell lysate of 1 L culture (OD_750_ ∼ 2) was split into multiple fractions. To detect the localization of endogenous IM30, the fractions of the lysate were mixed with Mg^2+^-containing HEPES buffer to achieve final concentrations of up to 20 mM Mg^2+^. As a control, HEPES buffer containing 20 mM EDTA was added to remove endogeneous Mg^2+^. The membrane fractions of the samples were isolated by ultracentrifugation (40000 rpm, 30 min, 4 °C). The supernatant was discarded, and the pellets were resuspended with buffer containing the respective amounts of Mg^2+^ and incubated 30 min. Finally, the pellets were resuspended to a final concentration of 1 mg/mL chlorophyll and 68% sucrose. The chlorophyll concentration was determined as decribed in (47) using methanol as a solvent. Membrane fractions containing 200 µg chlorophyll were loaded at the bottom of a linear 34 – 68% sucrose gradient. Cell fractions were separated by centrifugation at 40000 rpm for 6 h at 4 °C, and the gradients were immediately fractioned afterward. The TM fraction was identified by measuring the chlorophyll concentration of all gradient-fractions. The TM fractions were analyzed by SDS-PAGE and immunoblotting using an anti-IM30 antiserum (6). To compare the amounts of detected protein, the blots were subjected to densitometric analysis using ImageJ (46)

### Atomic Force Microscopy

All buffers and solutions were freshly prepared and filter sterilized (0.2 µM filter) before use. Freshly cleaved muscovite mica (12 mm diameter; Ted Pella Inc. grade V1) was mounted on a teflon disc (16 mm) and washed with 2*50 µL of adsorption buffer (20 mM HEPES pH 7.6, 20 mM Mg^2+^) to remove soluble ions from the mica surface. Afterward, the buffer was incubated for 5 min at room temperature on the mica disc. Thereafter, 50 µL liposome suspension (5 mg/mL DOPG) was added to the adsorption buffer. The mixture was incubated for 20 to 30 min at room temperature. Care was taken to let the substrate surface never run dry. The mica surface was washed carefully with 1 mL imaging buffer (20 mM HEPES w/o 10 mM Mg^2+^). A drop of 100 µL of imaging buffer was left on the mica. Then the sample was mounted under the AFM head. AFM measurements were carried out using a Nanowizard IV (JPK) in the QI mode (Quantitative imaging).

An uncoated AC240TS probe (OMCL-AC240TS: L=240 µm, W=40 µm, k=2 N/m; f=70kHz, Olympus) with a 7 nm tip was used for scanning. The formation of a lipid bilayer was checked by force curve measurements and indicated by the typical break-through shape of the force curve (48). A total concentration of ∼1.5 µM IM30 was achieved by adding small volumes (30 – 50 µL) directly on the lipid-coated mica surface mounted in the AFM. The sample was incubated at room temperature until the drift in the deflection signal decreased. Scanning was started with a minimal setpoint. Images were scanned with 256×256 or 512×512 px.

The resulting images were analyzed with GWYDDION version 2.56 (49). The measured height-signal images were leveled by removing a polynomial background, and scan rows were aligned using the median height. The images were cropped to the area of interest, where necessary and the offset was removed.

We analyzed the particle dimensions by marking them by a threshold, excluded membrane defects, particles spanning only one pixel, and subsequently exported the particle parameters via the analysis tool provided by GWYDDION.

### Quartz Crystal Microbalance

For QCM measurements, only degassed buffers were used. QCM chips were cleaned with 30 mM EDTA, 2% SDS followed by 1 M NaOH. Then, the chip was rinsed with water and dried with nitrogen. Prior to the measurement, the chip was treated with an ozone plasma cleaner for 20 s to remove any organic contaminations. 50 µL of a liposome suspension (1 mM lipid, 80% DOPC 20% DOPG (w/w)) was mixed with 450 µL 20 mM HEPES + 5 mM CaCl_2_. A SiO_2_-coated QCM chip (3T Analytik) was calibrated with the QCM device (3T Analytik). To produce an SLB on the SiO_2_ surface, the chip was washed with HEPES buffer. Next, 150 µL of the liposome suspension was pumped on the chip (60 µL/min). After completion of the liposome spreading (*i.e.* when the frequency shift reached a constant level of about 90 Hz and negligible damping), the chip was washed again with HEPES buffer or HEPES buffer + 10 mM Mg^2+^ (150+300 µL; 60 µL/min). To start the measurement, 150 µL IM30 wt (4.5 µM in HEPES buffer or in HEPES buffer + 10 mM Mg^2+^) was pumped on the SLB (60 µL/min) and binding of the protein was monitored over ∼3500 s.

## Data availability

The data supporting the findings of this study are available within the paper and its supplementary information files. All other relevant data are available upon reasonable request.

## Acknowledgments

This work was supported by the Max-Planck Graduate Center at the Max Planck institutes and the University of Mainz, as well as by DynaMem (state of Rhineland-Palatinate). We thank Annika Teresa Lehmann for support with the acquisition of the QCM data. Molecular graphics and analyses performed with UCSF ChimeraX, developed by the Resource for Biocomputing, Visualization, and Informatics at the University of California, San Francisco, with support from National Institutes of Health R01-GM129325 and the Office of Cyber Infrastructure and Computational Biology, National Institute of Allergy and Infectious Diseases.

## Abbreviations

IM30: Inner-membrane associated protein of 30 kDa
Vipp1: Vesicle inducing protein in plastids
PspA: Phage shock protein A
TM: Thylakoid membrane
CFP: Cyan fluorescent protein
GUV: Giant unilamellar vesicle
QCM: Quartz crystal microbalance
SLB: Solid supported bilayer
AFM: Atomic force microscopy
AMP: Antimicrobial peptide

